# The presence and impact of reference bias on population genomic studies of prehistoric human populations

**DOI:** 10.1101/487983

**Authors:** Torsten Günther, Carl Nettelblad

## Abstract

High quality reference genomes are an important resource in genomic research projects. A consequence is that DNA fragments carrying the reference allele will be more likely to map successfully, or receive higher quality scores. This reference bias can have effects on downstream population genomic analysis when heterozygous sites are falsely considered homozygous for the reference allele.

In palaeogenomic studies of human populations, mapping against the human reference genome is used to identify endogenous human sequences. Ancient DNA studies usually operate with low sequencing coverages and fragmentation of DNA molecules causes a large proportion of the sequenced fragments to be shorter than 50 bp – reducing the amount of accepted mismatches, and increasing the probability of multiple matching sites in the genome. These ancient DNA specific properties are potentially exacerbating the impact of reference bias on downstream analyses, especially since most studies of ancient human populations use pseudohaploid data, i.e. they randomly sample only one sequencing read per site.

We show that reference bias is pervasive in published ancient DNA sequence data of pre-historic humans with some differences between individual genomic regions. We illustrate that the strength of reference bias is negatively correlated with fragment length. Reference bias can cause differences in the results of downstream analyses such as population affinities, heterozygosity estimates and estimates of archaic ancestry. These spurious results highlight how important it is to be aware of these technical artifacts and that we need strategies to mitigate the effect. Therefore, we suggest some post-mapping filtering strategies to resolve reference bias which help to reduce its impact substantially.

## Introduction

The possibility to sequence whole genomes in a cost-efficient way has revolutionized the way how we do genetic and population genetic research. Annotated, high-quality reference genomes are a cornerstone for resequencing surveys which aim to study the genetic variation and demographic history of an entire species. Resequencing studies usually align the sequences of all studied individuals to a linear haploid reference sequence originating from a single individual or a mosaic of several individuals. In each site, this haploid sequence will only represent a single allele out of the entire genetic variation of the species. An inherent consequence is some degree of bias towards the alleles present in that reference sequence (“reference bias”). Sequencing reads carrying an alternative allele will naturally have mismatches in the alignment to the reference genome and consequently have lower mapping scores than reads carrying the same allele as the reference. This effect increases with genetic distance from the reference genome, which is of particular interest when using a reference genome from a related species for mapping (Shapiro and Hofreiter, 2014; Gopalakrishnan et al., 2017; Heintzman et al., 2017). Generally, reference bias can influence variant calling by missing alternative alleles or by wrongly calling heterozygous sites as homozygous reference (Bobo et al., 2016; Ros-Freixedes et al., 2018) which is known to influence estimates of heterozygosity and allele frequencies (Chen et al., 2012; Bryc et al., 2013; Brandt et al., 2015).

The field of palaeogenomics and the population genomic analysis of DNA obtained from hominin remains has led to a number of important insights and groundbreaking results in recent years, including admixture between different hominin groups, migrations of prehistoric humans and the evolution of different phenotypes (Günther and Jakobsson, 2016; Slatkin and Racimo, 2016; Nielsen et al., 2017; Dannemann and Racimo, 2018; Lazaridis, 2018; Skoglund and Mathieson, 2018). DNA preservation poses a major challenge for these studies, as fragmentation causes most authentic sequences to be shorter than 100 bp, and deamination damage increases the number of mismatches and can even mimic genetic variation at transition sites (Hofreiter et al., 2001; Brotherton et al., 2007; Briggs et al., 2007).

In addition to fragmentation and other post-mortem damages, low coverage data is a major limiting factor for ancient DNA studies. Coverages below 1x rarely permit calling diploid genotypes so a very common approach is to use “pseudo-haploid” data: at each known single nucleotide polymorphisms (SNP) site one sequencing read is picked at random (or following a majority rule) in order to represent a haploid genotype of that individual. This approach would not introduce bias if the reads were a random representation of the chromosomes carried by the individual. Reference bias, however, would introduce some skew towards the reference allele at heterozygous sites. These characteristics of ancient DNA and practices used in palaeogenomic studies make them particularly vulnerable to reference bias (Prüfer et al., 2010; Schubert et al., 2012). It has been shown that pseudo-haploid data can be more biased than imputed genotypes (Martiniano et al., 2017), and that reference bias and fragment length artifacts can interfere with phylogenetic classifications (Heintzman et al., 2017). Reference bias can influence downstream analyses if these are based on estimating allele frequencies in a population, or studying pairwise allele sharing between individuals and groups.

This study investigates the presence and impact of reference bias in studies of prehistoric human populations using genomic ancient DNA. We first illustrate its abundance in published data from ancient human and archaic hominins, and illustrate how it is influenced by standard data processing. We then show how reference bias can influence some basic population genetic analyses such as population affinities and heterozygosity. Finally, we discuss some possible data filtering strategies in order to mitigate reference bias in ancient DNA studies.

## Results

### Mapping quality filtering

We first investigate whether reference bias is present in published ancient DNA data. We restrict our analysis to known SNPs, as most population genomic analyses are using SNPs and the allele frequencies at those positions. In particular, we are only using transversion polymorphisms (to avoid the effect of post-mortem deaminations) and sites identified to be polymorphic in a world-wide set of modern human populations (Mallick et al., 2016). We investigate supposedly heterozygous sites (defined as sites covered by at least 10 reads with at least 25% representing the minor allele) in a set of published medium to high coverage human and hominin genomes (Table 1). We note that our approach does not include any rescaling of base qualities, as such approaches usually take the reference allele into account which may amplify reference bias.

**Table 1:** Information on the published medium to high coverage palaeogenomic and archaeogenomic data used in this study.

| Sample ID | (Partial) UDG treatment<sup>§</sup> | SNP capture | Average sequencing depth<sup>†</sup> | Reference |
| --- | --- | --- | --- | --- |
| Stuttgart | X |  | 15.8x | Lazaridis et al. (2014) |
| Loschbour | X |  | 17.7x | Lazaridis et al. (2014) |
| Ust-Ishim | X |  | 29.9x | Fu et al. (2014) |
| sf12 | X |  | 64.7x | Günther et al. (2018) |
| baa001 | X |  | 13.2x | Schlebusch et al. (2017) |
| Kotias |  |  | 12.5x | Jones et al. (2015) |
| Bichon |  |  | 15.4x | Jones et al. (2015) |
| ne1 |  |  | 18.5x | Gamba et al. (2014) |
| br2 |  |  | 15.4x | Gamba et al. (2014) |
| atp016 |  |  | 14.1x | Valdiosera et al. (2018) |
| Rathlin1 |  |  | 10.9x | Cassidy et al. (2015) |
| Ballynahatty |  |  | 10.7x | Cassidy et al. (2015) |
| I0054 | X | X | 2.4x | Mathieson et al. (2015) |
| I0103 | X | X | 2.4x | Mathieson et al. (2015) |
| I0118 | X | X | 1.7x | Mathieson et al. (2015) |
| I0172 | X | X | 3.4x | Mathieson et al. (2015) |
| I0408 | X | X | 1.7x | Mathieson et al. (2015) |
| I0412 | X | X | 1.9x | Mathieson et al. (2015) |
| I0585 | X | X | 2.4x | Mathieson et al. (2015) |
| AltaiNeandertal | X |  | 45.2x | Prüfer et al. (2014) |
| VindijaNeandertal | X |  | 25.9x | Prüfer et al. (2017) |
| Denisovan | X |  | 26.7x | Meyer et al. (2012) |

<sup>§</sup> Enzymatic repair of deamination damages
<sup>†</sup> at analyzed SGDP SNPs, using a minimum mapping quality of 30

At a heterozygous site, a DNA extract of an individual should contain the same number of reference and alternative fragments. We observe that after mapping to the human reference genome the average proportion of alternative alleles is lower than the expected 50 percent for all of the anatomically modern humans investigated (Figure 1), regardless of whether they represent SNP capture data, damage repaired libraries or standard shotgun sequencing (Table 1). As sequence fragments carrying the alternative allele will show an elevated number of mismatches to the reference genome, mapping quality seems a natural filter to avoid reference bias. Consistent with this expectation, we see a slightly stronger reference bias for stricter mapping quality filters. Lowering the mapping quality cutoff can have other detrimental effects, however, for example an enrichment of microbial contamination (Renaud et al., 2017) or sequences not uniquely mapping to a particular region of the genome. This is somewhat illustrated by the archaic genomes, two Neandertals and a Denisovan, which show - on average - a bias towards the alternative allele when no mapping quality filter is employed (Figure 1). This suggests that these more distant taxa carry variation in the genome which is not captured by the reference genome based on anatomically modern humans, in turn causing fragments originating in other parts of the genome to map at the investigated sites. As the qualities of the base calls have not been rescaled after mapping to the reference genome, we do not see an effect of different minimum base quality thresholds on reference bias (Supplementary Figure 1).

**Figure 1:**
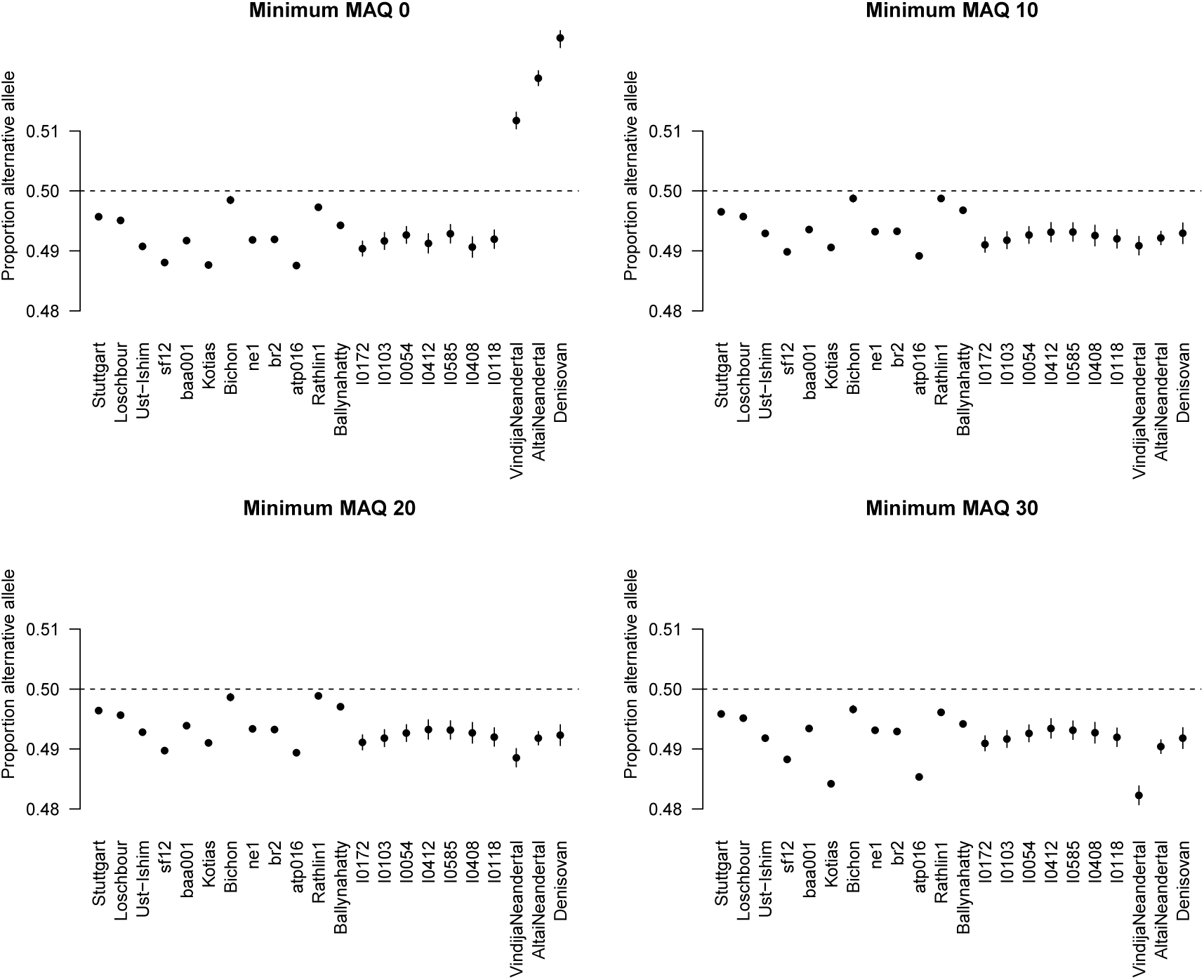
Reference bias in published genome-wide ancient DNA datasets for different minimum mapping quality thresholds. The plots show the average proportion of reads at heterozygous transversion sites (see Methods) representing the alternative allele. Error bars indicate two standard errors of the mean.

Investigating pairwise correlations between the proportion of alternative alleles at sites considered heterozygous in both individuals shows significantly positive correlations in most cases (Supplementary Table 1). This indicates that the strength of reference (and alternative) bias may differ regionally across the genome, so there could be an effect of sequence context and uniqueness of the specific sequences across the genome. The highest correlations are observed between samples from the same study or produced by the same institute suggesting that similar wet lab techniques also influence this effect.

### Distribution of bias

To investigate the distribution of reference bias instead of just averages as above, we modified original reads to carry opposite alleles at each SNP site and remapped them. We created a virtual read set for the Scandinavian Mesolithic hunter-gatherer sf12 (Günther et al., 2018) containing reads for all SNPs identified with mapping quality and base quality of at least 30 in the original mapping. No filter was placed on coverage, a SNP was included even if it was only covered by a single read. This joint read set of original and modified reads thus had perfectly balanced allele ratios for all SNPs. The full set was remapped, and SNPs were grouped based on the observed alternative allele fraction among all reads that again mapped to their respective SNPs with mapping quality of at least 30.

In total, 1,157,266 SNPs were analyzed. Out of these, 1,088,802 (94.08%) showed perfect allelic balance, with a proportion of alternative alleles of 0.5. About 100 SNPs were also affected by reads that mapped back with sufficient quality, but to a different genomic location. In these results, we only present proportions based on reads that map back to their original location from the first mapping round. The proportion of alternative alleles is summarized in Table 2. Notably, there is a subset of SNPs showing alternative, as opposed to reference, bias, and also a subset of SNPs where the bias is total, i.e. only one of the two alleles is ever mapped back successfully (within this dataset). The distribution across the genome of sites deviating from the balanced case is similar to the overall density of the SNPs used. Generally, all chromosomes and chromosomal regions are affected.

**Table 2:** Proportion of alternative alleles when mapping back original reads and virtual opposite allele reads for the sf12 individual.

| Proportion of alternative alleles | # SNPs | Percentage |
| --- | --- | --- |
| 0 | 11497 | 0.99% |
| (0, 0.4) | 12962 | 1.12% |
| [0.4, 0.5) | 37262 | 3.22% |
| 0.5 | 1088802 | 94.08% |
| (0.5, 0.6] | 4172 | 0.36% |
| (0.6, 1) | 1413 | 0.12% |
| 1 | 1158 | 0.10% |

### The influence of fragment length

Most mapping strategies set the number of allowed mismatches relative to the length of the sequenced fragment. Therefore, shorter fragments might show a stronger reference bias than long fragments. To investigate this, we used the 57x genome generated for the Scandinavian Mesolithic hunter-gatherer sf12 (Günther et al., 2018) and partitioned the data into fragment length bins. The large amount of data allows us to still have a sufficient number of SNPs covered at 10x or more for each of the length bins.

Somewhat expectedly, shorter fragments display a stronger reference bias than longer sequences (Figure 2A). Generally, fragment length might be a main driver of reference bias across all samples as the mode of each individual’s fragment size distribution is highly correlated with the average proportion of alternative alleles at heterozygous sites (Pearson’s *r* = 0.67, *p* = 0.0006; Figure 2B). This also has an effect on the proportion of sites considered heterozygous among all sites analyzed which can be seen as a relative measure for the individual’s heterozygosity (Figure 2C). In fact, different fragment length bins of the same individual produce heterozygosity estimates that do not overlap in their 95% confidence interval (Figure 2C). This represents a general limitation for estimating heterozygosity from ancient DNA data which may to some degree explain the generally low diversity estimates for many prehistoric groups (e.g. Skoglund et al., 2014; Kousathanas et al., 2017; Scheib et al., 2018). The potential of obtaining significantly different estimates for the same population genetic statistic may also have enormous effects on other downstream analyses such as population affinities and population structure.

**Figure 2:**
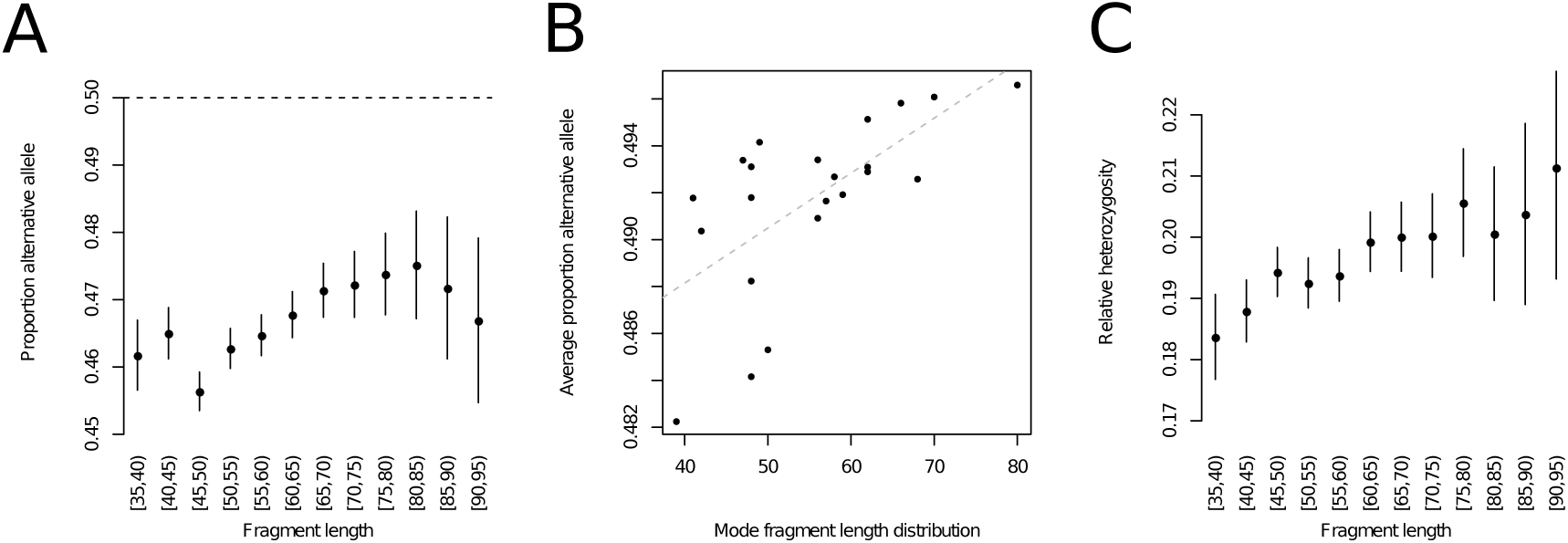
Connection between fragment length and reference bias. (A) Proportion of alternative allele for different fragment length bins in the high coverage individual sf12. (B) Correlation between average proportion of alternative alleles and the mode of the fragment size distribution across all investigated individuals. (C) Proportion of heterozygous sites among all sites with sufficient coverage for different fragment length bins in the high coverage individual sf12. All error bars indicate two standard errors.

### Impact on measures of population affinity

In order to investigate the influence of reference bias on population affinities, we calculated different combinations of *D* statistics of the form *D*(*Chimp, X*; *Y, Z*), where *X* is a modern human population, and *Y* and *Z* are two different treatments of the same individual sf12. Therefore, the expectation for *D* is 0, but differences in reference bias between *Y* and *Z* could lead to spurious allele sharing between population *X* and a deviation from 0. Negative values of *D* indicate more allele sharing of *X* with *Y* while positive values indicate an excess of shared alleles between *X* and *Z*. The populations *X* were grouped by continental origin and we calculated the statistics separately for whole genome shotgun data (SGDP, Mallick et al., 2016) and genotyped populations (HO, Lazaridis et al., 2014).

We use four different versions of genotypes for sf12. First, we compare pseudo-haploid calls (random allele per site with minimum mapping and base quality of 30) to diploid genotype calls (Figure 3A and C). This comparison assumes that the diploid calls are less affected by reference bias as slight deviations from a 50/50-ratio at heterozygous sites should be tolerated by a diploid genotype caller but random sampling would be biased towards the reference allele. This is supported by the *D* statistic *D*(*chimp, reference*; *sf* 12_*hapl, sf* 12_*dipl*) < 0 (*Z* = −13.5), indicating more allele sharing between the reference and the pseudo-haploid calls. For this illustration, we are using diploid genotype calls from *GATK* as we are only looking at the variation at known SNP sites. We note that genotype callers specifically developed for ancient DNA (Link et al., 2017; Zhou et al., 2017; Prüfer, 2018) are preferable when calling novel variants from ancient DNA data as they incorporate post-mortem damage and other ancient DNA specific properties. Second, we compare randomly sampled reads of different fragment length categories (Figure 3B and D) as longer (75-80 bp) fragments should exhibit less reference bias than short (35-40 bp) fragments (see above), which is supported by the *D* statistic *D*(*chimp, reference*; *sf* 12_*short, sf* 12_*long*) < 0 (*Z* = −20.6), indicating more allele sharing between the reference and pseudo-haploid calls from short fragments.

**Figure 3:**
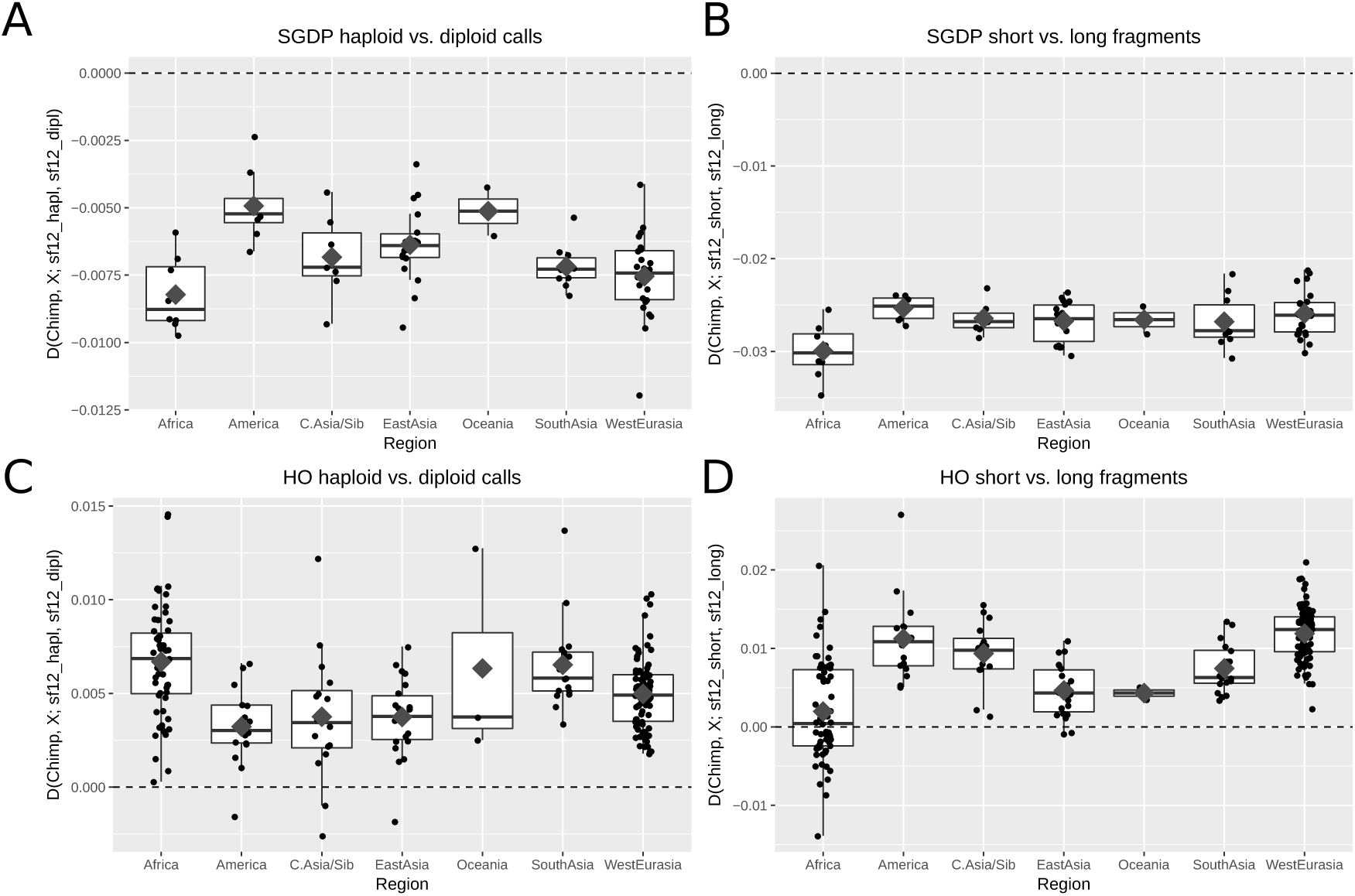
*D* statistics testing the affinity between different modern populations (*X*) and two different treatments of the high coverage individual sf12. The basis for these comparisons are the modern sequence data of the SGDP panel (A and B) or genotype data from the HO panel (C and D). Comparisons are done between pseudo-haploid and diploid calls for sf12 (A and C), and between pseudo-haploid calls from short (35-40 bp) or long (75-80 bp) fragments (B and D). The x axis represents the geographic origin of population *X*, diamonds show the mean for each continental group.

In general, we observe a deviation from zero in most cases highlighting the effect of reference bias on these statistics (Figure 3). Surprisingly, the directions of this bias differ between the HO data and the SGDP data, which suggests that different reference data sets are also affected by reference bias at different degrees. This represents a potential batch effect which also needs to be considered when merging different reference data sets. Affinities to populations of different geographic origin vary in their sensitivity to reference bias but little general trends are observable. Western Eurasian populations show a strong deviation from 0 in all tests. Notably, African populations show the strongest deviation in the short versus long comparison in the SGDP data set while they exhibit almost no bias in the same comparison using the HO data. As the biases do not seem to show a consistent tendency, we cannot directly conclude that recent ancient DNA papers have been systematically biased in some direction. The shifts appear to be dataset and test specific so some results could still be driven by spurious affinities due to reference bias.

The human reference genome sequence is a mosaic of the genomes of different individuals. The geographic origin of the specific segments should have an impact on the population genetic affinities as the reference allele will more likely be found in specific geographic regions. We obtained information on the local ancestry of the human reference genome from Green et al. (2010). According to this estimate 15.6 % of the reference genome can be assigned to African, 5.0 % to East Asian and 30.0 % to European origin while the origin for 49.4 % is uncertain. We re-calculate *D* statistics for the different parts of the genome separately, restricting the analysis to the SGDP data. The impact of reference bias differs between the different ancestries (Figure 4). Generally, reference bias is weakest for reference segments of African origin. Notably, African populations show the strongest deviations from 0 in this case. Sequences mapping to the European segments of the reference show a strong reference bias with slight differences between continental populations. Reference bias at the East Asian segments of the reference genome seems intermediate but the *D* statistics also show large variation which may be due to the only small proportion of the reference genome that could confidently be assigned to an East Asian origin (Green et al., 2010).

**Figure 4:**
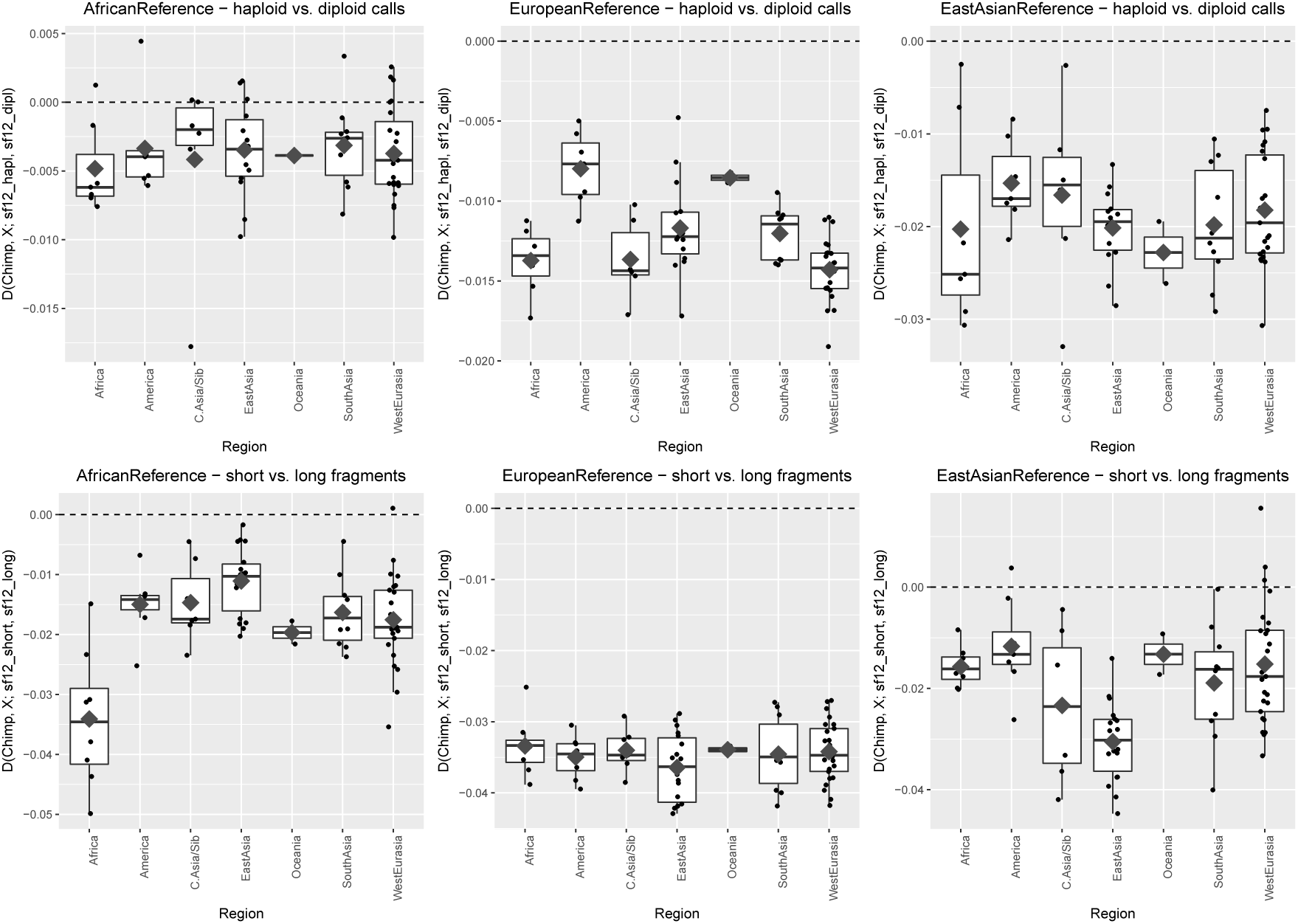
*D* statistics similar to Figure 3 for different parts of the reference genome depending on their geographic origin (Green et al., 2010). The x axis represents the geographic origin of population *X*, diamonds show the mean for each continental group.

Finally, we explore whether reference bias can affect estimates of archaic ancestry. We estimate the Neandertal ancestry proportion in sf12 as done by Prüfer et al. (2017):

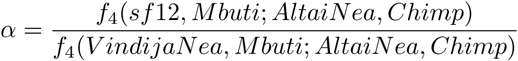

We use eight different combinations of diploid and pseudo-haploid calls for sf12 as well as the two Neandertals in this statistic (Table 3). The 95% confidence intervals of all estimates overlap but point estimates differ by up to 1.25% when using all pseudo-haploid versus all diploid calls. The African segments of the reference genome yield the lowest point estimates (as low as 1.42%) – some of these estimates are not even significantly different from 0. These differences highlight some of the sensitivities of *f*_4_-ratios not just to the choice of reference populations (Petr et al., 2018) but also to technical artifacts.

**Table 3:**
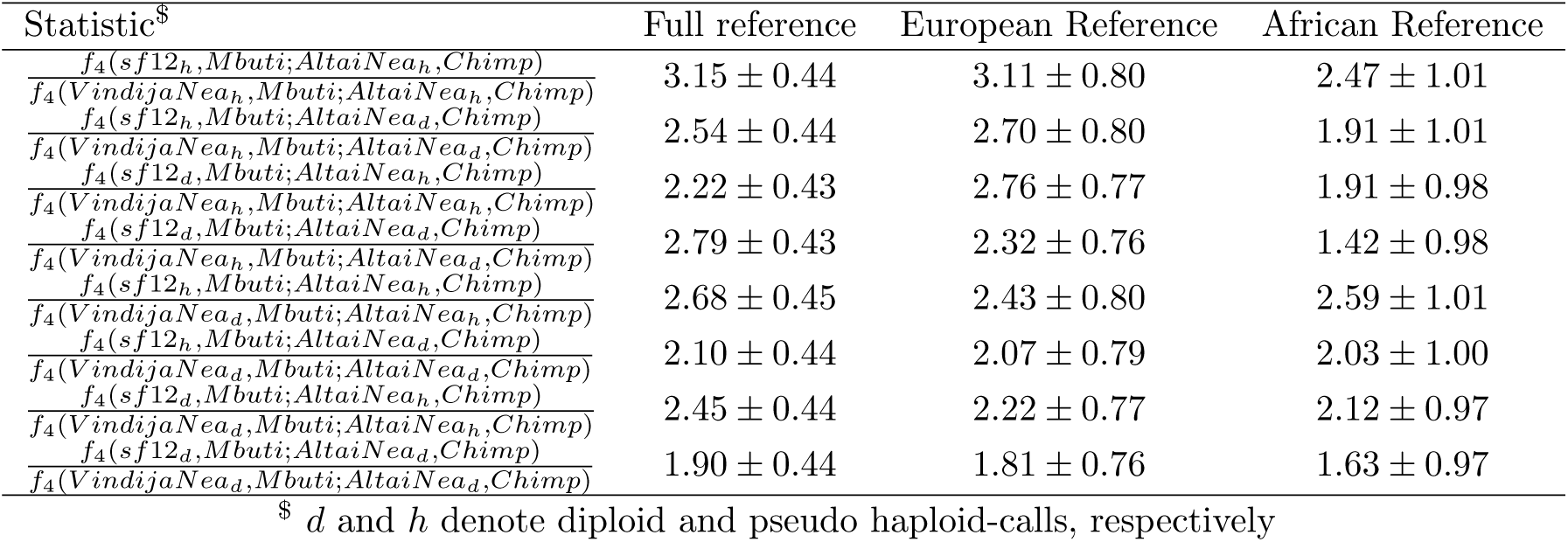
Percentage of Neandertal ancestry (and standard errors) in sf12 using diploid and pseudohaploid calls and different subsets of the human reference genome. Parts of the genome of East Asian origin were excluded due to their small total size.

### Potential data filtering strategies

After establishing the abundance and potential effect of reference bias, we investigated two simple post-mapping filtering approaches to mitigate reference bias. The two agents involved in the process are the reference genome and the sequence fragments or reads. We investigated 1,407,340 of the SGDP transversion set of sites with at least 200 bp distance between two neighboring SNPs. First, we modified reads that successfully mapped to a SNP site with a match of the reference allele to carry the alternative allele. These modified reads were re-mapped to the reference genome and they passed the filtering if they still mapped to the same position of the genome with no indels. Second, we prepared a modified version of the reference genome which carried a third base (neither the reference base nor the known alternative allele) at all 1,407,340 sites. A similar approach has been used to study ultra-short fragments in sequence data from archaic hominins (de Filippo et al., 2018). All reads originally mapping to the SNP sites were re-mapped to this modified reference genome, and again only reads that mapped to the same location and without indels passed the filtering. Finally, we used both filters on the same BAM file. All scripts used for filtering can be found at https://bitbucket.org/tguenther/refbias/

The filtering approaches increase the average proportion of the alternative allele at homozygous sites (Figure 5A). Mapping to a modified reference genome shows a slightly better improvement than using modified reads, while combining both yields the best results in most cases. A small number of samples shows a 50/50-ratio after filtering but most are still significantly below that ratio. This is not surprising as the filtering is only applied to reads that have previously mapped to a single reference genome so the data before filtering does not represent a 50/50-ratio, and removing some reference allele reads cannot completely account for the non-reference reads lost earlier. This is most evident in the data from Mathieson et al. (2015) which was only available as mapped reads after running *bwa* (Li and Durbin, 2009) with lower maximum edit distance parameters (-n 0.04) than our pipeline which does not leave much room for improvement after filtering. Another possible reason for deviation from a 50/50-ratio at heterozygous sites could be low levels of modern contamination which may lead to a slight over-representation of the reference allele before mapping (Prüfer et al., 2014; Racimo et al., 2016; Prüfer, 2018). Comparing the outcome of the filters to different fragment length categories shows a similar pattern: the bias is decreased but some length categories still display differences in their relative heterozygosity (Figure 5B).

**Figure 5:**
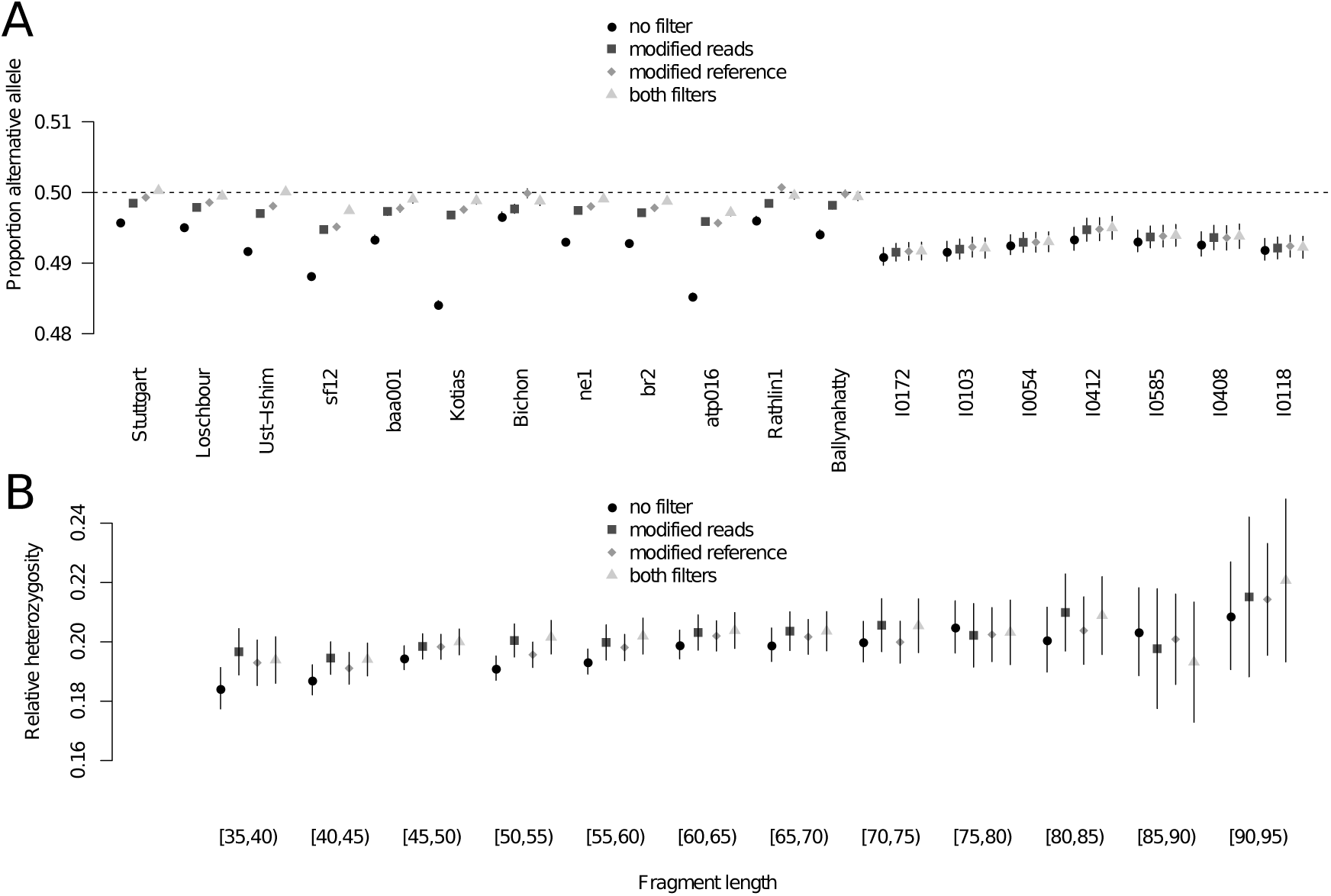
Comparison of different post-mapping filtering strategies for high coverage bam files from anatomically modern humans employing mapping and base quality filters of 30. (A) Average proportion of the alternative allele for the comparison between no additional filters (see also Figure 1), remapping of reads carrying the reference allele modified to carry the alternative allele (modified reads), remapping against a modified reference carrying a third allele at the SNP sites, and both filters together. (B) Influence of filtering on measures of heterozygosity for different fragment sizes in sf12. Error bars indicate two standard errors.

## Discussion

Systematic biases are problematic in all types of quantitative research, and it is therefore important to be aware of them and alleviate or avoid their effects as much as possible. Different systematic biases in next-generation sequencing data have been investigated before (Prüfer et al., 2010; Ross et al., 2013; Bobo et al., 2016; Ros-Freixedes et al., 2018), and it is known parameters such as sequencing depth can influence population genomic estimates (Crawford and Lazzaro, 2012; Fumagalli, 2013; Korneliussen et al., 2013). Differences in sequencing strategies (e.g. read length) and bioinformatic processing have been shown to generate batch effects and dramatically affect downstream analyses (Leek et al., 2010; Leigh et al., 2018; Shafer et al., 2016; Mafessoni et al., 2018). Another well known bias in population genetics is ascertainment bias which arises when the studied variants were ascertained in selected populations only, and can substantially impact measurements of heterozygosity and related methods (Albrechtsen et al., 2010). The research community is aware of these potential issues and they are avoided by filtering strategies, standardizing bioinformatic pipelines, including controls and accounting for systematic biases in downstream analysis.

The common use of randomly sampled alleles and pseudo-haploid data in palaeogenomic research can exacerbate the effect of reference bias compared to diploid genotype calls obtained from medium to high coverage data. We show that reference bias is able to lead to significant differences between estimates of population genetic parameters (heterozygosity), overestimated levels of archaic ancestry as well as to cause spurious affinities to certain populations. Mixing different mapping parameters or minimum fragment lengths in the same study should generally be avoided. Additionally, strong differences of fragment size distributions between different individuals may cause spurious affinities due to reference bias. Many estimates from low coverage data are generally noisy, but studies show increasing sample sizes and amounts of data which means that subtle biases become of increasing importance in the future. Notably, the bias for the whole genome (Figure 3) seems less extreme than some of the results for ancestry-specific segments (Figure 4) suggesting that the mosaic nature of the human reference genome may reduce the bias to some degree as different regions will be biased in different directions. In this respect the human reference genome is different from many other species where the reference genome is derived from a single individual which would increase the potential impact of reference bias on population genetic analysis in other systems.

Our analysis does not directly indicate a strong direct impact of different wet lab procedures on the observed average degree of reference bias. We caution, however, that such an effect may exists as indicated by the correlations between samples processed in the same lab (Supplementary Table 1). Different library preparation techniques produce different length distributions since some approaches are directly targeting shorter fragments which will have an impact on mapping. Furthermore, the SNP capture approaches used to generate the data we analyzed uses one bait per allele minimizing reference bias before sequencing. Most whole genome or exome capture approaches, however, are using baits designed from a single individual which should introduce a pre-mapping bias towards the allele carried by that person (Quail et al., 2008; Heinrich et al., 2012; Meynert et al., 2013; Lindo et al., 2016). Finally, contamination from another person should tend to introduce the major allele which is likely the reference allele in most cases – a process that will also increase reference bias before mapping (Prüfer et al., 2014; Racimo et al., 2016; Prüfer, 2018).

Our analysis of the distribution of reference bias across the genome for the sf12 individual has several repercussions. First, most reads are neutral to changing the allele to its opposing counterpart. This leads to a possible alternative filtering strategy. In cases where a pre-defined set of variants is acceptable, a quality control should be performed on the study level to filter out SNPs which correspond to reads that do not survive this alternative mapping. The exact details of such a filter will, again, be dependent on the expected length and degradation of the reads. Another important observation is that reference bias does not operate alone. There is also a weaker, but very clear, signal of alternative allele bias, affecting roughly 0.6% of the total SNPs. In addition, both reference and alternative bias can sometimes be very strong on the level of individual SNPs. Even in a dataset with an overall proportion of alternative reads close to 0.5 in heterozygous sites overall, subsets of SNPs might perform very differently, again possibly confusing deeper forms of analysis that do not only consider genome-wide metrics – for example selection scans or analysis of loci involved in certain traits.

We show, that filtering steps can reduce but not completely eliminate reference bias at SNPs after mapping. To fully prevent reference bias, alternative mapping strategies would be needed or filtering strategies would have to be developed for all raw data which is not always published. Furthermore, these proposed filters require a pre-defined set of variants used for downstream analysis and are not suitable for calling novel variants from ancient DNA data. The latter, however, will generally be only restricted to high quality and high coverage samples. A recently developed genotype caller for ancient DNA data estimates reference bias from the data and uses the estimate as a parameter for variant calling (Prüfer, 2018), which seems to work well for samples sequenced to coverages of 15x or higher. One could use the filtering steps tested by us in a similar manner to estimate what proportion of reads in a library are affected by reference bias which could later be used to estimate genotype likelihoods (Nielsen et al., 2011; Wang et al., 2013). As reference bias is somewhat predictable and detectable, this offers opportunities to account for it in downstream analyses (e.g. Bryc et al., 2013; Wu et al., 2017).

Alternative mapping strategies such as mapping against genome graphs (Paten et al., 2017; Garrison et al., 2018) or multiple reference genomes simultaneously (Schneeberger et al., 2009) could be able to eliminate reference bias already in the mapping step. These approaches are not broadly established in human genomics yet but their development has huge potential with regard to reference bias. Such approaches could also lead to an increase in the total amount of authentic data that can be obtained from a library while post-mapping filters will reduce the amount of data used for downstream analyses (between 2 and 10 % in our cases). In addition to filtering data and standardizing bioinformatic pipelines for all samples used in a study (both published data and newly sequenced), we propose simulations as a potential control. Specific ancient DNA simulation suites (Renaud et al., 2017) provide the opportunity to simulate data exactly matching fragment size and damage patterns of empirical ancient DNA data so one can use them to study if observed patterns may be driven by reference bias alone.

The present study focused mainly on humans but the effect of reference bias extends to other species as well. The slight alternative bias in archaic hominins and the different population affinities depending on the geographic origin of the reference genome illustrate that increasing evolutionary distance can exacerbate reference bias or even cause systematic alternative bias at some sites. This suggests that mapping against a reference genome of a related species (in the absence of a reference genome for the species in focus) may impact downstream analyses as well (Green et al., 2010; Schubert et al., 2012; Shapiro and Hofreiter, 2014; Gopalakrishnan et al., 2017), but the population genetic bias may be weaker as the reference genome employed usually represents an outgroup of equal distance to all individuals in the studied species.

## Conclusion

Our analysis highlights that reference bias is pervasive in ancient DNA data used to study prehistoric populations. While the strength of the effect differs between applications and data set, it is clear that reference bias has the potential to create spurious results in population genomic analyses. Furthermore, even when the overall presence of bias is limited, it is important to assess whether subsets of variants are prone to strong systematic bias, including the possible presence of alternative bias.

We are entering a time where sample sizes in ancient DNA studies reach one hundred and beyond, while the questions focus on more and more detailed patterns and subtle differences. At the same time, sampling starts to involve older remains and remains from more challenging environments – both of which are usually associated with poor preservation and shorter fragments. Therefore it seems crucial to avoid reference bias or other biases such as batch effects or ascertainment biases as much as possible, and to develop and apply computational strategies to mitigate the impact of these issues.

## Materials and Methods

### Data sets and bioinformatic processing

We selected medium to high coverage data from 22 different individuals representing data generated by different research groups with different wet lab strategies, covering different geographic regions and time periods (Table 1). For anatomically modern human samples, we tried to use data as raw as possible but some publications only provided the data after mapping and filtering. The general pipeline for these samples was identical to previous studies (Günther et al., 2015, 2018). Reads were mapped to the 1000 genomes version of the human reference genome hg19 using *bwa* (Li and Durbin, 2009) with non-default parameters -l 16500 -n 0.01 -o 2. Subsequently, PCR duplicates and fragments shorter than 35 bp were filtered (Kircher, 2012).

We restricted our analysis to a set of known transversion variants to avoid an effect of postmortem damage. We selected 107,404 transversions from the Human Origins panel (Patterson et al., 2012; Lazaridis et al., 2014) as well as 1,693,337 transversions which were at at least 5% allele frequency in the public data of the Simons Genome Diversity Project (SGDP, Mallick et al., 2016). To detect reference bias, we are looking at supposedly heterozygous sites where one would expect reads to map in a 50/50-ratio on average if no bias existed. We define a heterozygous site as a SNP for which we observe at least ten reads with between 25 to 75% of those representing the alternative allele. These reads are assessed using *samtools mpileup* (version 1.5, Li et al., 2009) employing the -B option to turn off base quality rescaling.

For the high coverage genome of sf12 (Günther et al., 2018) as well as the high coverage archaic genomes (Meyer et al., 2012; Prüfer et al., 2014, 2017) we also generated diploid genotype calls following the pipeline described in Günther et al. (2018). Briefly, base qualities of all Ts in the first five base pairs of each read as well as all As in the last five base pairs were set to 2. *Picard* version 1.118 (Broad Institute, 2016) was used to add read groups to the files followed by indel realignment with *GATK* 3.5.0 (McKenna et al., 2010) based on reference indels identified in phase 1 of the 1000 genomes project (Auton et al., 2015). Finally, diploid genotypes were called with *GATK* ’s UnifiedGenotyper employing the parameters -stand_call_conf 50.0, -stand_emit_conf 50.0, -mbq 30, -contamination 0.02 and –output_mode EMIT_ALL_SITES using dbSNP version 142 as known SNPs. Genotype calls not flagged as low quality calls at investigated SNP sites were extracted from the VCF files using *vcftools* (Danecek et al., 2011).

### Population genetic tests

In order to investigate the population genetic effect of reference bias, we calculated *D* and *f* statistics (Patterson et al., 2012). These statistics are based on pairwise allele sharing, so they should be sensitive to spurious allele sharing due to reference bias. *D* statistics were calculated with *popstats* (Skoglund et al., 2015), *f*4 ratios were calculated *ADMIXTOOLS* (Patterson et al., 2012), and standard errors were calculated employing a weighted block jackknife with a block size of 5 Mbp. We used the chimpanzee reference genome as an outgroup.

### Data Access

All scripts used for filtering can be found at https://bitbucket.org/tguenther/refbias/

## Supporting information

## Acknowledgements

We are grateful to Arielle Munters, Federico Sanchez, Mattias Jakobsson, and other members of the Human Evolution research program for discussions and comments as well as the attendees of various early presentations on this topic for their input and encouragement to turn it into a manuscript. We also thank Arielle Munters for initial data processing and Shop Mallick for sharing the local ancestry information for the human reference genome. The computations were performed on resources provided by SNIC through Uppsala Multidisciplinary Center for Advanced Computational Science (UPPMAX) under projects sllstore2017087, uppstore2018139, SNIC 2018/8-106 and 2018/8-239. TG was supported by grants from the Swedish Research Council Vetenskapsrådet and The Royal Physiographic Society of Lund (Nilsson-Ehle Endowments), as well as a Knut och Alice Wallenbergs Stiftelse grant to Mattias Jakobsson. CN was supported by a grant from the Swedish Research Council for Environment, Agricultural Sciences and Spatial Planning (FORMAS).

## References

Albrechtsen, A., Nielsen, F. C., and Nielsen, R., 2010. Ascertainment Biases in SNP Chips Affect Measures of Population Divergence. Molecular Biology and Evolution, 27(11):2534–2547.

Auton, A., Abecasis, G. R., Altshuler, D. M., Durbin, R. M., Abecasis, G. R., Bentley, D. R., Chakravarti, A., Clark, A. G., Donnelly, P., Eichler, E. E., et al., 2015. A global reference for human genetic variation. Nature, 526(7571):68–74.

Bobo, D., Lipatov, M., Rodriguez-Flores, J. L., Auton, A., and Henn, B. M., 2016. False Negatives Are a Significant Feature of Next Generation Sequencing Callsets. bioRxiv, :066043.

Brandt, D. Y. C., Aguiar, V. R. C., Bitarello, B. D., Nunes, K., Goudet, J., and Meyer, D., 2015. Mapping Bias Overestimates Reference Allele Frequencies at the HLA Genes in the 1000 Genomes Project Phase I Data. G3: Genes, Genomes, Genetics, 5(5):931–941.

Briggs, A. W., Stenzel, U., Johnson, P. L., Green, R. E., Kelso, J., Prüfer, K., Meyer, M., Krause, J., Ronan, M. T., Lachmann, M., et al., 2007. Patterns of damage in genomic DNA sequences from a Neandertal. Proceedings of the National Academy of Sciences, 104(37):14616–14621.

Broad Institute, 2016. Picard tools. https://broadinstitute.github.io/picard/,.

Brotherton, P., Endicott, P., Sanchez, J. J., Beaumont, M., Barnett, R., Austin, J., and Cooper, A., 2007. Novel high-resolution characterization of ancient DNA reveals C> U-type base modification events as the sole cause of post mortem miscoding lesions. Nucleic acids research, 35(17):5717–5728.

Bryc, K., Patterson, N. J., and Reich, D., 2013. A Novel Approach to Estimating Heterozygosity from Low-Coverage Genome Sequence. Genetics, genetics.113.154500.

Cassidy, L. M., Martiniano, R., Murphy, E. M., Teasdale, M. D., Mallory, J., Hartwell, B., and Bradley, D. G., 2015. Neolithic and Bronze Age migration to Ireland and establishment of the insular Atlantic genome. Proceedings of the National Academy of Sciences, :1–6.

Chen, X., Listman, J. B., Slack, F. J., Gelernter, J., and Zhao, H., 2012. Biases and Errors on Allele Frequency Estimation and Disease Association Tests of Next-Generation Sequencing of Pooled Samples. Genetic Epidemiology, 36(6):549–560.

Crawford, J. E. and Lazzaro, B. P., 2012. Assessing the accuracy and power of population genetic inference from low-pass next-generation sequencing data. Frontiers in Genetics, 3:66.

Danecek, P., Auton, A., Abecasis, G., Albers, C. A., Banks, E., DePristo, M. A., Handsaker, R. E., Lunter, G., Marth, G. T., Sherry, S. T., et al., 2011. The variant call format and VCFtools. Bioinformatics (Oxford, England), 27(15):2156–2158.

Dannemann, M. and Racimo, F., 2018. Something old, something borrowed: admixture and adaptation in human evolution. Current Opinion in Genetics & Development, 53:1–8.

de Filippo, C., Meyer, M., and Prüfer, K., 2018. Quantifying and reducing spurious alignments for the analysis of ultra-short ancient DNA sequences. BMC Biology, 16(1):121.

Fu, Q., Li, H., Moorjani, P., Jay, F., Slepchenko, S. M., Bondarev, A. A., Johnson, P. L. F., Aximu-Petri, A., Prüfer, K., de Filippo, C., et al., 2014. Genome sequence of a 45,000-year-old modern human from western Siberia. Nature, 514(7523):445–449.

Fumagalli, M., 2013. Assessing the Effect of Sequencing Depth and Sample Size in Population Genetics Inferences. PLOS ONE, 8(11):e79667.

Gamba, C., Jones, E. R., Teasdale, M. D., McLaughlin, R. L., Gonzalez-Fortes, G., Mattiangeli, V., Domboroczki, L., Kövári, I., Pap, I., Anders, A., et al., 2014. Genome flux and stasis in a five millennium transect of European prehistory. Nature Communications, 5:5257.

Garrison, E., Sirén, J., Novak, A. M., Hickey, G., Eizenga, J. M., Dawson, E. T., Jones, W., Garg, S., Markello, C., Lin, M. F., et al., 2018. Variation graph toolkit improves read mapping by representing genetic variation in the reference. Nature Biotechnology,.

Gopalakrishnan, S., Samaniego Castruita, J. A., Sinding, M.-H. S., Kuderna, L. F. K., Räikkönen, J., Petersen, B., Sicheritz-Ponten, T., Larson, G., Orlando, L., Marques-Bonet, T., et al., 2017. The wolf reference genome sequence (Canis lupus lupus) and its implications for Canis spp. population genomics. BMC Genomics, 18:495.

Green, R. E., Krause, J., Briggs, A. W., Maricic, T., Stenzel, U., Kircher, M., Patterson, N., Li, H., Zhai, W., Fritz, M. H.-Y., et al., 2010. A draft sequence of the Neandertal genome. Science (New York, N.Y.), 328(5979):710–22.

Günther, T. and Jakobsson, M., 2016. Genes mirror migrations and cultures in prehistoric Europe-a population genomic perspective. Current Opinion in Genetics & Development, 41:115–123.

Günther, T., Malmström, H., Svensson, E. M., Omrak, A., Sánchez-Quinto, F., Kılınç, G. M., Krzewińska, M., Eriksson, G., Fraser, M., Edlund, H., et al., 2018. Population genomics of Mesolithic Scandinavia: Investigating early postglacial migration routes and high-latitude adaptation. PLoS biology, 16(1):e2003703.

Günther, T., Valdiosera, C., Malmström, H., Ureña, I., Rodriguez-Varela, R., Sverrisdottir, O. O., Daskalaki, E. A., Skoglund, P., Naidoo, T., Svensson, E. M., et al., 2015. Ancient genomes link early farmers from Atapuerca in Spain to modern-day Basques. Proceedings of the National Academy of Sciences of the United States of America, 112(38):11917–11922.

Heinrich, V., Stange, J., Dickhaus, T., Imkeller, P., Krüger, U., Bauer, S., Mundlos, S., Robinson, P. N., Hecht, J., and Krawitz, P. M., et al., 2012. The allele distribution in next-generation sequencing data sets is accurately described as the result of a stochastic branching process. Nucleic Acids Research, 40(6):2426–2431.

Heintzman, P. D., Zazula, G. D., MacPhee, R. D., Scott, E., Cahill, J. A., McHorse, B. K., Kapp, J. D., Stiller, M., Wooller, M. J., Orlando, L., et al., 2017. A new genus of horse from Pleistocene North America. eLife, 6.

Hofreiter, M., Jaenicke, V., Serre, D., Haeseler, A. v., and Pääbo, S., 2001. DNA sequences from multiple amplifications reveal artifacts induced by cytosine deamination in ancient DNA. Nucleic acids research, 29(23):4793–4799.

Jones, E. R., Gonzalez-Fortes, G., Connell, S., Siska, V., Eriksson, A., Martiniano, R., McLaughlin, R. L., Gallego Llorente, M., Cassidy, L. M., Gamba, C., et al., 2015. Upper Palaeolithic genomes reveal deep roots of modern Eurasians. Nature communications, 6:8912.

Kircher, M., 2012. Analysis of high-throughput ancient DNA sequencing data. volume 840, pages 197–228.

Korneliussen, T. S., Moltke, I., Albrechtsen, A., and Nielsen, R., 2013. Calculation of Tajima’s D and other neutrality test statistics from low depth next-generation sequencing data. BMC bioinformatics, 14:289.

Kousathanas, A., Leuenberger, C., Link, V., Sell, C., Burger, J., and Wegmann, D., 2017. Inferring Heterozygosity from Ancient and Low Coverage Genomes. Genetics, 205(1):317–332.

Lazaridis, I., 2018. The evolutionary history of human populations in Europe. Current Opinion in Genetics & Development, 53:21–27.

Lazaridis, I., Patterson, N., Mittnik, A., Renaud, G., Mallick, S., Kirsanow, K., Sudmant, P. H., Schraiber, J. G., Castellano, S., Lipson, M., et al., 2014. Ancient human genomes suggest three ancestral populations for present-day Europeans. Nature, 513(7518):409–413.

Leek, J. T., Scharpf, R. B., Bravo, H. C., Simcha, D., Langmead, B., Johnson, W. E., Geman, D., Baggerly, K., and Irizarry, R. A., 2010. Tackling the widespread and critical impact of batch effects in high-throughput data. Nature Reviews Genetics, 11(10):733–739.

Leigh, D. M., Lischer, H. E. L., Grossen, C., and Keller, L. F., 2018. Batch effects in a multiyear sequencing study: False biological trends due to changes in read lengths. Molecular Ecology Resources, 0(0).

Li, H. and Durbin, R., 2009. Fast and accurate short read alignment with Burrows-Wheeler transform. Bioinformatics (Oxford, England), 25(14):1754–1760.

Li, H., Handsaker, B., Wysoker, A., Fennell, T., Ruan, J., Homer, N., Marth, G., Abecasis, G., Durbin, R., and 1000 Genome Project Data Processing Subgroup, et al., 2009. The Sequence Alignment/Map format and SAMtools. Bioinformatics (Oxford, England), 25(16):2078–2079.

Lindo, J., Huerta-Sánchez, E., Nakagome, S., Rasmussen, M., Petzelt, B., Mitchell, J., Cybulski, J. S., Willerslev, E., DeGiorgio, M., and Malhi, R. S., et al., 2016. A time transect of exomes from a Native American population before and after European contact. Nature Communications, 7:13175.

Link, V., Kousathanas, A., Veeramah, K., Sell, C., Scheu, A., and Wegmann, D., 2017. ATLAS: analysis tools for low-depth and ancient samples. bioRxiv, :105346.

Mafessoni, F., Prasad, R. B., Groop, L., Hansson, O., Prüfer, K., and McLysaght, A., 2018. Turning vice into virtue: Using Batch-Effects to Detect Errors in Large Genomic Datasets. Genome Biology and Evolution,.

Mallick, S., Li, H., Lipson, M., Mathieson, I., Gymrek, M., Racimo, F., Zhao, M., Chennagiri, N., Nordenfelt, S., Tandon, A., et al., 2016. The Simons Genome Diversity Project: 300 genomes from 142 diverse populations. Nature, 538(7624):201–206.

Martiniano, R., Cassidy, L. M., Ó’Maoldúin, R., McLaughlin, R., Silva, N. M., Manco, L., Fidalgo, D., Pereira, T., Coelho, M. J., Serra, M., et al., 2017. The population genomics of archaeological transition in west Iberia: Investigation of ancient substructure using imputation and haplotypebased methods. PLoS genetics, 13(7):e1006852.

Mathieson, I., Lazaridis, I., Rohland, N., Mallick, S., Patterson, N., Roodenberg, S. A., Harney, E., Stewardson, K., Fernandes, D., Novak, M., et al., 2015. Genome-wide patterns of selection in 230 ancient Eurasians. Nature, 528(7583):499–503.

McKenna, A., Hanna, M., Banks, E., Sivachenko, A., Cibulskis, K., Kernytsky, A., Garimella, K., Altshuler, D., Gabriel, S., Daly, M., et al., 2010. The Genome Analysis Toolkit: A MapReduce framework for analyzing next-generation DNA sequencing data. Genome Research, 20(9):1297–1303.

Meyer, M., Kircher, M., Gansauge, M.-T., Li, H., Racimo, F., Mallick, S., Schraiber, J. G., Jay, F., Prüfer, K., de Filippo, C., et al., 2012. A high-coverage genome sequence from an archaic Denisovan individual. Science (New York, N.Y.), 338(6104):222–226.

Meynert, A. M., Bicknell, L. S., Hurles, M. E., Jackson, A. P., and Taylor, M. S., 2013. Quantifying single nucleotide variant detection sensitivity in exome sequencing. BMC Bioinformatics, 14:195.

Nielsen, R., Akey, J. M., Jakobsson, M., Pritchard, J. K., Tishkoff, S., and Willerslev, E., 2017. Tracing the peopling of the world through genomics. Nature, 541(7637):302–310.

Nielsen, R., Paul, J. S., Albrechtsen, A., and Song, Y. S., 2011. Genotype and SNP calling from next-generation sequencing data. Nature Reviews Genetics, 12(6):443.

Paten, B., Novak, A. M., Eizenga, J. M., and Garrison, E., 2017. Genome graphs and the evolution of genome inference. Genome Research, 27(5):665–676.

Patterson, N., Moorjani, P., Luo, Y., Mallick, S., Rohland, N., Zhan, Y., Genschoreck, T., Webster, T., and Reich, D., 2012. Ancient Admixture in Human History. Genetics, 192(3):1065–1093.

Petr, M., Pääbo, S., Kelso, J., and Vernot, B., 2018. The limits of long-term selection against Neandertal introgression. bioRxiv, :362566.

Prüfer, K., 2018. snpAD: An ancient DNA genotype caller. Bioinformatics,.

Prüfer, K., Filippo, C. d., Grote, S., Mafessoni, F., Korlević, P., Hajdinjak, M., Vernot, B., Skov, L., Hsieh, P., Peyrégne, S., et al., 2017. A high-coverage Neandertal genome from Vindija Cave in Croatia. Science, 358(6363):655–658.

Prüfer, K., Racimo, F., Patterson, N., Jay, F., Sankararaman, S., Sawyer, S., Heinze, A., Renaud, G., Sudmant, P. H., de Filippo, C., et al., 2014. The complete genome sequence of a Neanderthal from the Altai Mountains. Nature, 505(7481):43–9.

Prüfer, K., Stenzel, U., Hofreiter, M., Pääbo, S., Kelso, J., and Green, R. E., 2010. Computational challenges in the analysis of ancient DNA. Genome Biology, 11:R47.

Quail, M. A., Kozarewa, I., Smith, F., Scally, A., Stephens, P. J., Durbin, R., Swerdlow, H., and Turner, D. J., 2008. A large genome center’s improvements to the Illumina sequencing system. Nature Methods, 5(12):1005–1010.

Racimo, F., Renaud, G., and Slatkin, M., 2016. Joint estimation of contamination, error and demography for nuclear DNA from ancient humans. PLoS genetics, 12(4):e1005972.

Renaud, G., Hanghøj, K., Willerslev, E., and Orlando, L., 2017. gargammel: a sequence simulator for ancient DNA. Bioinformatics, 33(4):577–579.

Ros-Freixedes, R., Battagin, M., Johnsson, M., Gorjanc, G., Mileham, A. J., Rounsley, S. D., and Hickey, J. M., 2018. Impact of index hopping and bias towards the reference allele on accuracy of genotype calls from low-coverage sequencing. Genetics Selection Evolution, 50(1).

Ross, M. G., Russ, C., Costello, M., Hollinger, A., Lennon, N. J., Hegarty, R., Nusbaum, C., and Jaffe, D. B., 2013. Characterizing and measuring bias in sequence data. Genome Biology, 14(5):R51.

Scheib, C. L., Li, H., Desai, T., Link, V., Kendall, C., Dewar, G., Griffith, P. W., Mörseburg, A., Johnson, J. R., Potter, A., et al., 2018. Ancient human parallel lineages within North America contributed to a coastal expansion. Science, 360(6392):1024–1027.

Schlebusch, C. M., Malmström, H., Günther, T., Sjödin, P., Coutinho, A., Edlund, H., Munters, A. R., Vicente, M., Steyn, M., Soodyall, H., et al., 2017. Southern African ancient genomes estimate modern human divergence to 350,000 to 260,000 years ago. Science (New York, N.Y.),

Schneeberger, K., Hagmann, J., Ossowski, S., Warthmann, N., Gesing, S., Kohlbacher, O., and Weigel, D., 2009. Simultaneous alignment of short reads against multiple genomes. Genome Biology, 10(9):R98.

Schubert, M., Ginolhac, A., Lindgreen, S., Thompson, J. F., AL-Rasheid, K. A., Willerslev, E., Krogh, A., and Orlando, L., 2012. Improving ancient DNA read mapping against modern reference genomes. BMC Genomics, 13:178.

Shafer, A. B. A., Peart, C. R., Tusso, S., Maayan, I., Brelsford, A., Wheat, C. W., and Wolf, J. B. W., 2016. Bioinformatic processing of RAD-seq data dramatically impacts downstream population genetic inference. Methods in Ecology and Evolution, 8(8):907–917.

Shapiro, B. and Hofreiter, M., 2014. A paleogenomic perspective on evolution and gene function: new insights from ancient DNA. Science (New York, N.Y.), 343(6169):1236573.

Skoglund, P., Mallick, S., Bortolini, M. C., Chennagiri, N., Hünemeier, T., Petzl-Erler, M. L., Salzano, F. M., Patterson, N., and Reich, D., 2015. Genetic evidence for two founding populations of the Americas. Nature, 525(7567):104–108.

Skoglund, P., Malmstrom, H., Omrak, A., Raghavan, M., Valdiosera, C., Gunther, T., Hall, P., Tambets, K., Parik, J., Sjogren, K.-G., et al., 2014. Genomic Diversity and Admixture Differs for Stone-Age Scandinavian Foragers and Farmers. Science, 344(6185):747–750.

Skoglund, P. and Mathieson, I., 2018. Ancient Human Genomics: The First Decade. Annual Review of Genomics and Human Genetics, 19(1):ull.

Slatkin, M. and Racimo, F., 2016. Ancient DNA and human history. Proceedings of the National Academy of Sciences of the United States of America, 113(23):6380–6387.

Valdiosera, C., Günther, T., Vera-Rodríguez, J. C., Ureña, I., Iriarte, E., Rodríguez-Varela, R., Simões, L. G., Martínez-Sánchez, R. M., Svensson, E. M., Malmström, H., et al., 2018. Four millennia of Iberian biomolecular prehistory illustrate the impact of prehistoric migrations at the far end of Eurasia. Proceedings of the National Academy of Sciences, :201717762.

Wang, Y., Lu, J., Yu, J., Gibbs, R. A., and Yu, F., 2013. An integrative variant analysis pipeline for accurate genotype/haplotype inference in population NGS data. Genome Research, 23(5):833–842.

Wu, S. H., Schwartz, R. S., Winter, D. J., Conrad, D. F., and Cartwright, R. A., 2017. Estimating error models for whole genome sequencing using mixtures of Dirichlet-multinomial distributions. Bioinformatics, 33(15):2322–2329.

Zhou, B., Wen, S., Wang, L., Jin, L., Li, H., and Zhang, H., 2017. AntCaller: an accurate variant caller incorporating ancient DNA damage. Molecular Genetics and Genomics, 292(6):1419–1430.

