## Supplementary figures and images for "The presence and impact of reference bias on population genomic studies of prehistoric human populations"

### Supplementary file 1

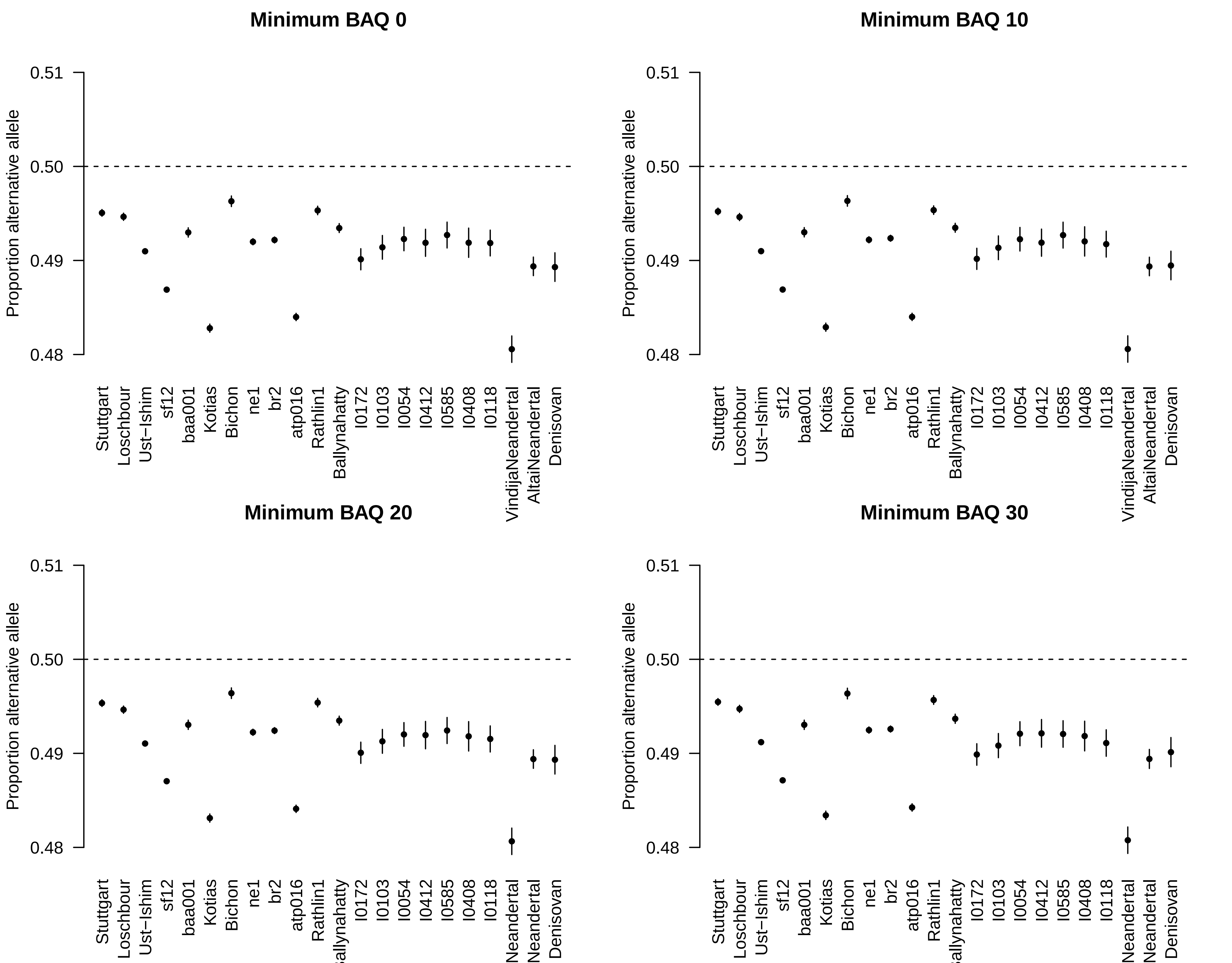
